# Decoding mouse behavior to explain single-trial decisions and their relationship with neural activity

**DOI:** 10.1101/567479

**Authors:** Yves Weissenberger, Andrew J. King, Johannes C. Dahmen

**Affiliations:** Department of Physiology, Anatomy and Genetics, University of Oxford, Oxford, UK.

## Abstract

Models of behavior typically focus on sparse measurements of motor output over long timescales, limiting their ability to explain momentary decisions or neural activity. We developed data-driven models relating experimental variables to videos of behavior. Applied to mouse operant behavior, they revealed behavioral encoding of cognitive variables. Model-based decoding of videos yielded an accurate account of single-trial behavior in terms of the relationship between cognition, motor output and cortical activity.

## Main Text

Advances in neural recording technologies have enabled activity to be measured from thousands of neurons simultaneously^1,2^. By eliminating the need for averaging activity across trials, these methods are providing unprecedented insights into neural function. But to fully realize their promise, we also require similarly comprehensive descriptions of behavior that can be used to bridge the gap between neural activity and function.

However, even in highly-controlled experimental settings, such as during a sensory decision-making task, quantitative descriptions of behavioral variability remain elusive^3,4^. Analyses of session-level choice-statistics have shown that decisions are influenced by a variety of factors^5,6^. Nevertheless, it remains extremely challenging to identify the factors underlying single-trial decisions from currently available behavioral readouts. This severely limits the functional interpretation of brain activity, which often relies on such behavioral readouts to link neural activity to cognitive processes.

The interpretation of neural activity is further complicated by correlations between experimental variables (e.g. cognitive variables or environmental stimuli) and motor output. Indeed, such correlations can confound the neural encoding of an experimental variable like a decision with the encoding of the associated motor output, i.e. the enactment of the decision.

One approach to overcoming these issues is the detailed quantitative study of behavior^4^. Classical approaches^7^ focus on simple measures (e.g. aggregate choice-statistics) that are easy to relate back to experimental variables. However, these measures lack the capacity or temporal resolution that is required to robustly link neural activity to the computations underpinning trial-by-trial behavior. Although recent approaches have begun to address these shortcomings by performing unsupervised decompositions of detailed behavioral measurements^8,9^, their output can be difficult to relate to experimental variables, thereby limiting their scope.

We sought a novel and generally applicable approach to the challenge of quantifying behavior which combines the strengths of previous methods. We took a data-driven approach and developed statistical models of dense behavioral measurements. Our objective was to find representations of behavior that can account for an animal’s motor output whilst remaining easily relatable to cognitive and stimulus-related variables. Crucially, we attempted to find such representations directly in the data, without *a priori* knowledge. In doing so, we aimed to extract a comprehensive and interpetable account of behavior that can support detailed analysis of neural activity.

We analyzed video data from head-fixed mice (n = 11 sessions from 6 mice) performing a sound detection task (**Fig. 1a**), and used variational autoencoders, which are Bayesian latent-variable models (LVM)^10,11^, as a starting point for modelling animals’ motor output. The aim of the model was to find low-dimensional representations of the video data that enable frame-by-frame reconstructions at pixel-level resolution (**Fig. 1b i**).

**Figure 1.**
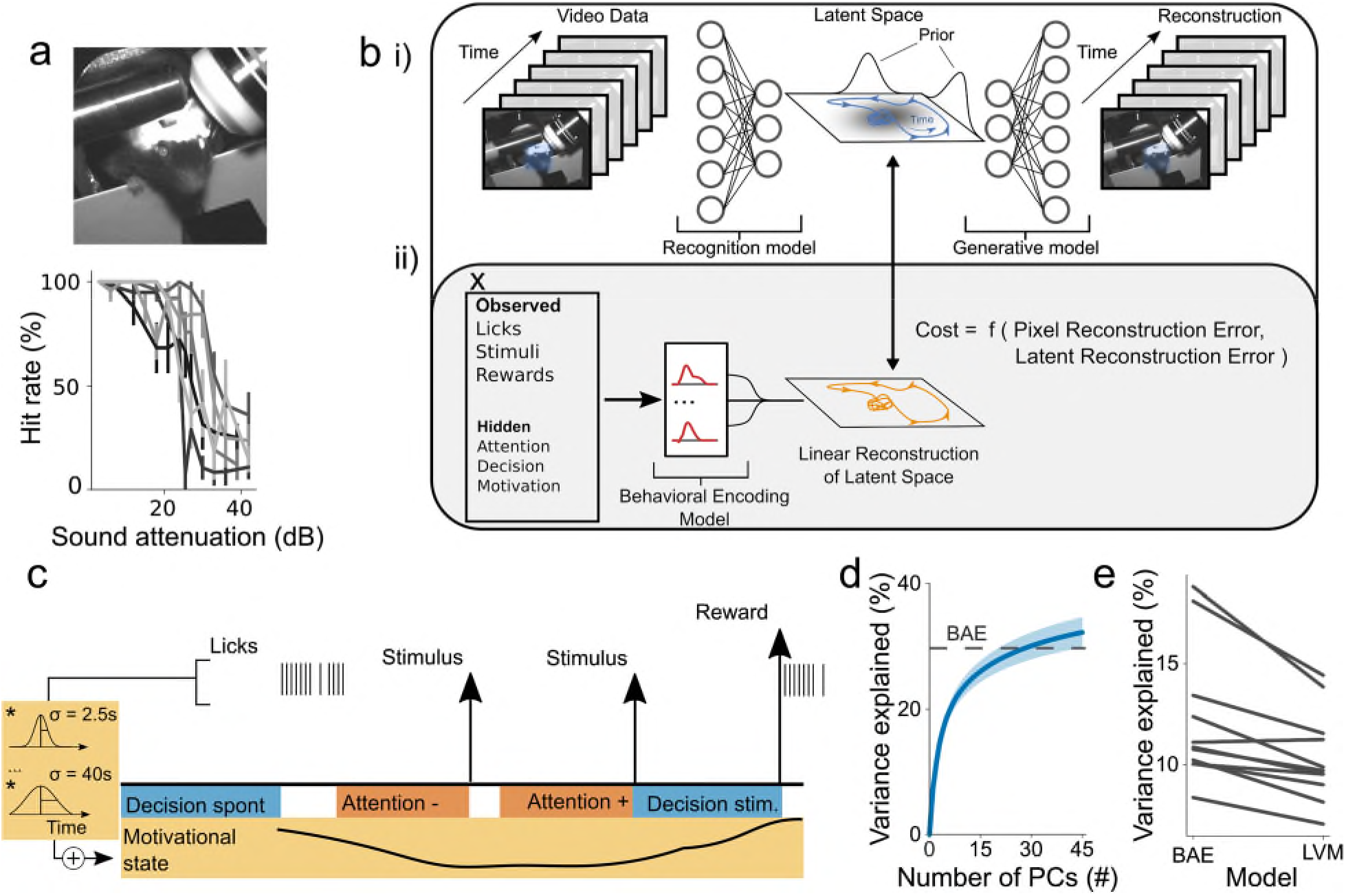
Model Structure and performance. **a) *(top)*** Image of a mouse in the experimental setup. ***(bottom)*** Example psychometric functions (±95% binomial confidence intervals) illustrating performance in the sound detection task (each curve depicts performance of one mouse in a single session; all curves are from different mice). **b)** Schematic of the LVM and BAE. **(i)** The LVM is parameterized by two sequential deep neural networks. The first network parameterizes a recognition model that maps from video data to a low-dimensional latent space. The second network parameterizes a generative model which maps from the latent space back into pixel space and reconstructs the video data. **(ii)** The BAE encompasses the LVM and a behavioral encoding model that maps experimental variables into an approximation of the latent space. This is used to encourage latent representations to be linearly predictable from experimental variables **x** by an additional penalty term, which structures representations in the latent space. **c)** Schematic illustrating the definition of hidden variables (see *Methods).* Briefly, an animal was considered attentive on a given trial if the stimulus was of low intensity and the trial was a hit-trial. It was considered inattentive on a given trial if the stimulus was of low intensity and the trial was a miss-trial. An animal was considered to engage in ‘stimulus-driven’ licking if a stimulus occurred in a 540-ms window preceding the onset of a lick bout; otherwise the licking was considered to be ‘spontaneous’. A high lick rate was interpreted to be indicative of reward seeking and, thus, a state of high motivation. Motivational state regressors were created by convolving licks with a series of Gaussian filters that were fitted individually and then summed. Relative timescales across elements of the figure are not to scale. **d)** Performance of the BAE (dashed line; latent states were inferred using the recognition model) compared with a principal component analysis (PCA) based reconstruction (mean ±2 s.e.m) as a function of number of PCs. Here, BAE reconstructions used the recognition model. **e)** Comparison of the LVM and the BAE’s ability to reconstruct videos using the behavioral encoding model (paired-sample t-test; *p* = 4.1 · 10^−4^).

Models of behavior are useful only to the extent that they can be related to experimental variables, such as an animal’s decisions or the underlying neural activity. We therefore formalized the notion of relatability as linear predictability from these variables. This yielded a novel model, which we refer to as a behavioral autoencoder (BAE), the cost function of which is augmented with an additional penalty term. This term encourages learning a representation of behavior that is explicable in terms of *a priori* defined variables of interest (**Fig. 1b ii**) (see *Methods).* We then fitted this model to videos acquired during task performance.

The sound detection task provided a rich set of observed and hidden variables (**Fig. 1c**), which may explain momentary variations in animals’ motor output. We therefore used both sets of variables (henceforth referred to collectively as experimental variables) to augment the model’s cost function.

To assess the model’s performance, we quantified the reconstruction quality and capacity of the experimental variables to explain behavioral latent states. Qualitative and quantitative analyses revealed accurate reconstruction of the video data (mean r^2^ = 30%, s.e.m = 3%) (**Supplementary Fig. 1, Supplementary Video 1**). Quantitatively, a 10-dimensional BAE outperformed optimal linear methods, which required three-fold greater dimensionality to account for the same variance (**Fig. 1d, Supplementary Fig. 2a**). Importantly, learned representations were highly interpretable, as assayed by measuring their predictability from experimental variables (**Supplementary Fig. 2b**). Furthermore, augmentation of the cost function in the BAE significantly improved this predictability over that provided by the LVM (**Fig. 1e, Supplementary Fig. 2b**). Together, these findings suggest that the model learned comprehensive and interpretable representations of the animals’ behavior.

We then asked which experimental variables were encoded (i.e. expressed) in the animals’ behavior by quantifying the capacity of individual variables to explain behavioral latent states. Although we found that all variables are encoded in behavior (**Fig. 2a**), this may arise simply because many of them are correlated. We therefore quantified the effect of excluding subsets of regression parameters, relating to a single experimental variable, on model-fit quality (see *Methods).* This revealed that only a subset of variables uniquely accounted for variance in the data (**Fig. 2b**). Time into session accounted for most variance, reflecting the fact that the animals’ resting posture gradually changed over the course of the session. Additionally, we consistently found that the animals’ motivational state (operationalized as a smoothed lick time series, **Fig. 1c**; see *Methods*) was explicitly encoded in behavior (**Supplementary Fig. 3a,b**). By contrast, we found no evidence that trial-by-trial variations in attention or stimulus presentation were expressed in behavior (**Fig. 2a,b, Supplementary Fig. 3c**). The latter result suggests that the animals’ behavioral response to the stimulus is largely embodied by its decision to lick.

**Figure 2.**
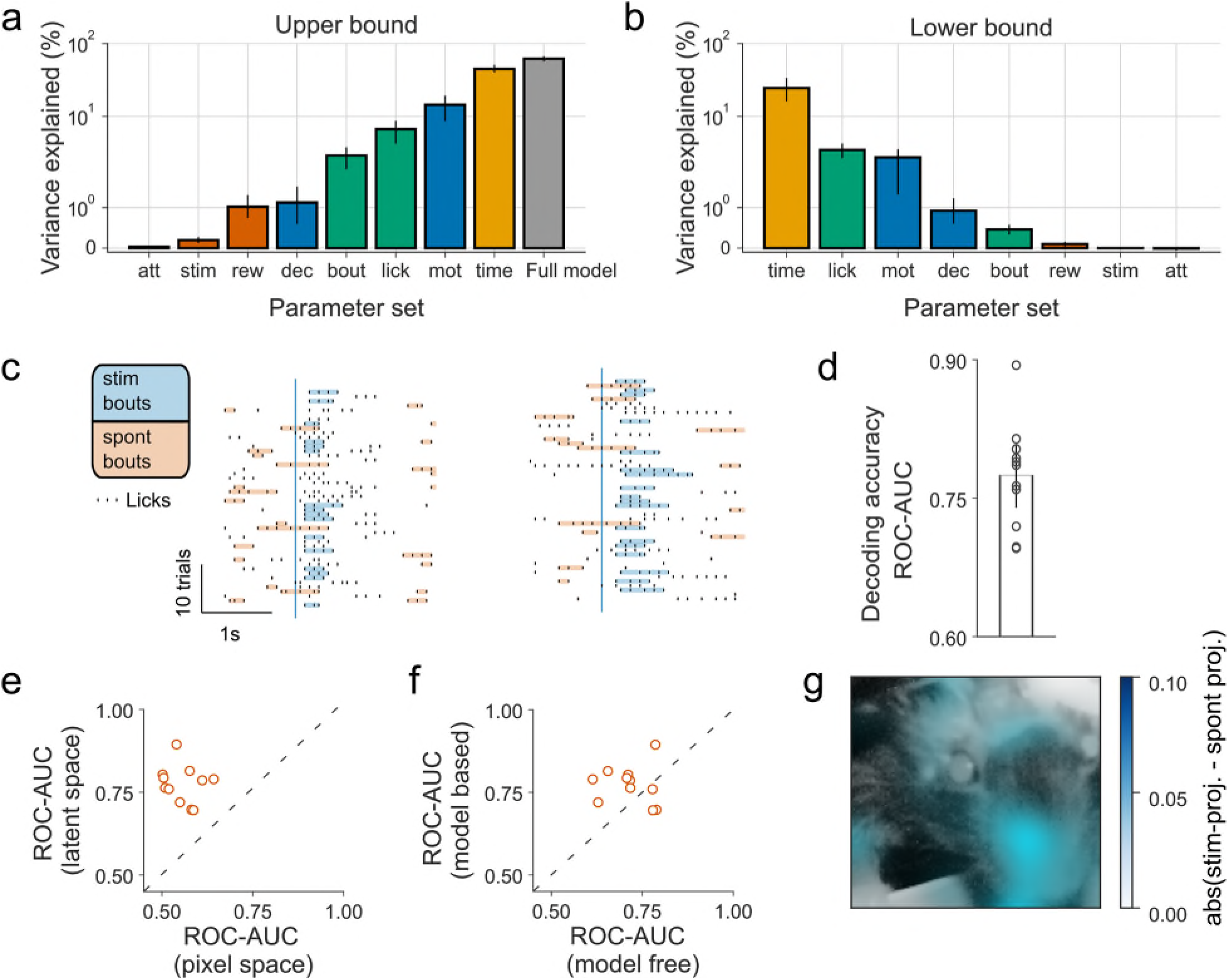
Encoding and decoding behavior. **a)** Estimation of upper bounds on extent of encoding by only regressing parameter sets belonging to a single variable. Variables are sorted according to their ability to predict latent states. **b)** Estimation of lower bound on extent of encoding by removing regressors relating to a single variable, one at a time, and subtracting cross-validated r^2^ for full model performance from r^2^ for models with individual components removed. Error bars show bootstrapped 95% confidence intervals. **c)** Excerpts from two example sessions showing lick-bouts defined as either stimulus-driven (blue) or spontaneous (orange) depending on their timing relative to the stimulus onset (blue vertical line). **d)** Decoding of intention (i.e. classification of bout type) by inverting behavioral encoding models reveals accurate decoding (mean ROC-AUC = 0.78; s.e.m = 0.01). Error-bars show ± 2 s.e.m. Circles are individual data-points. **e)** Decoding in latent space is more accurate than decoding in pixel space (paired samples t-test; *p* =3.9 · 10^−6^). **f)** Model-based decoding performs better than model-free (SVM) decoding (paired samples t-test; *p* = 0.0086). **g)** Difference between the BAE’s estimate of a stimulus and a spontaneous bout overlayed on an image of a mouse. Estimates were created by projecting linear predictions of stimulus-driven and spontaneous bouts into pixel space. In this case, informative pixels are clustered around the snout. (att=attention; stim=stimulus presentation; rew=reward delivery; dec=decision basis (spontaneous vs stimulus-driven licking); bout=lick-bout initiation; mot=motivational state)

Given the importance of single-trial analyses in decision-making paradigms^12,13^, we next investigated the behavioral correlates of decision-making processes. The non-zero false alarm rates observed in our data suggest that multiple processes drive mouse licking. We therefore sought to test whether distinct causes of licking (i.e. spontaneous vs. stimulus-driven) were differentially encoded in behavior (**Fig. 1c, Fig. 2a,b**). To do so, we attempted to decode the causes of licking on a lick-by-lick basis.

We grouped licks into bouts (**Fig. 2c, Supplementary Fig. 4**) and selected a counterbalanced set (see *Methods*) of stimulus-driven (fast response times on trials with loud stimuli) and spontaneous (outside of the peri-stimulus period) lick-bouts. We then decoded (i.e. predicted) the causes of these bouts using the latent states within the ~500 ms preceding the first lick of each bout. Previous work has demonstrated that the inversion of encoding models offers a powerful and parsimonious approach to decoding^14,15^. We therefore constructed model-based decoders based on the inversion of the behavioral-encoding models (**Fig. 1b**). Consistent with results from the encoding perspective, we were able to decode, on a bout-by-bout basis, whether a stimulus preceded a bout or not (**Fig. 2d**). Thus, the animals’ behavior preceding a lick bout allowed us to infer whether a stimulus drove that bout.

Further analysis demonstrated that decoding accuracy was higher in the latent-space than in pixel-space (**Fig. 2e**) and that model-based decoding out-performed comparable model-free support vector machines (SVM) (**Fig. 2f**). Importantly, decoding is unlikely to be driven by motor preparation (**Supplementary Fig. 5a-d**). Finally, the generative capabilities of the BAE enabled us to project linear approximations of stimulus-driven and spontaneous lick bouts back into pixel space. This visual account of the basis of their classification revealed that idiosyncratic behaviors associated with lick bouts formed the basis for classification (**Fig. 2g, Supplementary Videos 2,3**).

Model-based decoding thus offers a data-driven alternative to *a priori* analysis of behavior. In doing so, it both provides a way of automatically identifying behavioral correlates of experimental variables and a means of classifying behavior based on these correlates. In turn, this yields an interpretable account of momentary behavior that can readily be employed to improve our understanding of neural activity.

To demonstrate this, we sought to explicitly benchmark model-based and *a priori* classifications of trial-by-trial decisions against neural activity. Previous work has demonstrated that behavioral choice correlates with the activity of neurons in primary auditory cortex (A1)^16–18^. We reasoned that by comparing the behavioral categorization of bout-by-bout intent with neural activity, we would be able to compare the two classification approaches.

We therefore performed two-photon calcium imaging of excitatory layer 2/3 neurons in A1 of three mice (**Fig. 3a-c**). To assess whether neural activity covaries with behavioral choice, we computed choice probabilities^12^ (CPs), and identified a subpopulation of L2/3 neurons with significant CPs (**Fig. 3d,e; Supplementary Fig. 6**). CPs calculated by comparing hit-trials and miss-trials were both significantly correlated with (**Fig. 3f**) and not systematically different from (**Supplementary Fig. 7a**) those calculated by comparing hit-trials with level-matched hit-trials in which animals responded prematurely (i.e. with a latency of <120 ms, which is faster than mouse reaction times). These results argue that CPs reflected sensorimotor coupling, rather than licking or reward consumption, and were thus used as a benchmark measure of behavioral classification.

**Figure 3.**
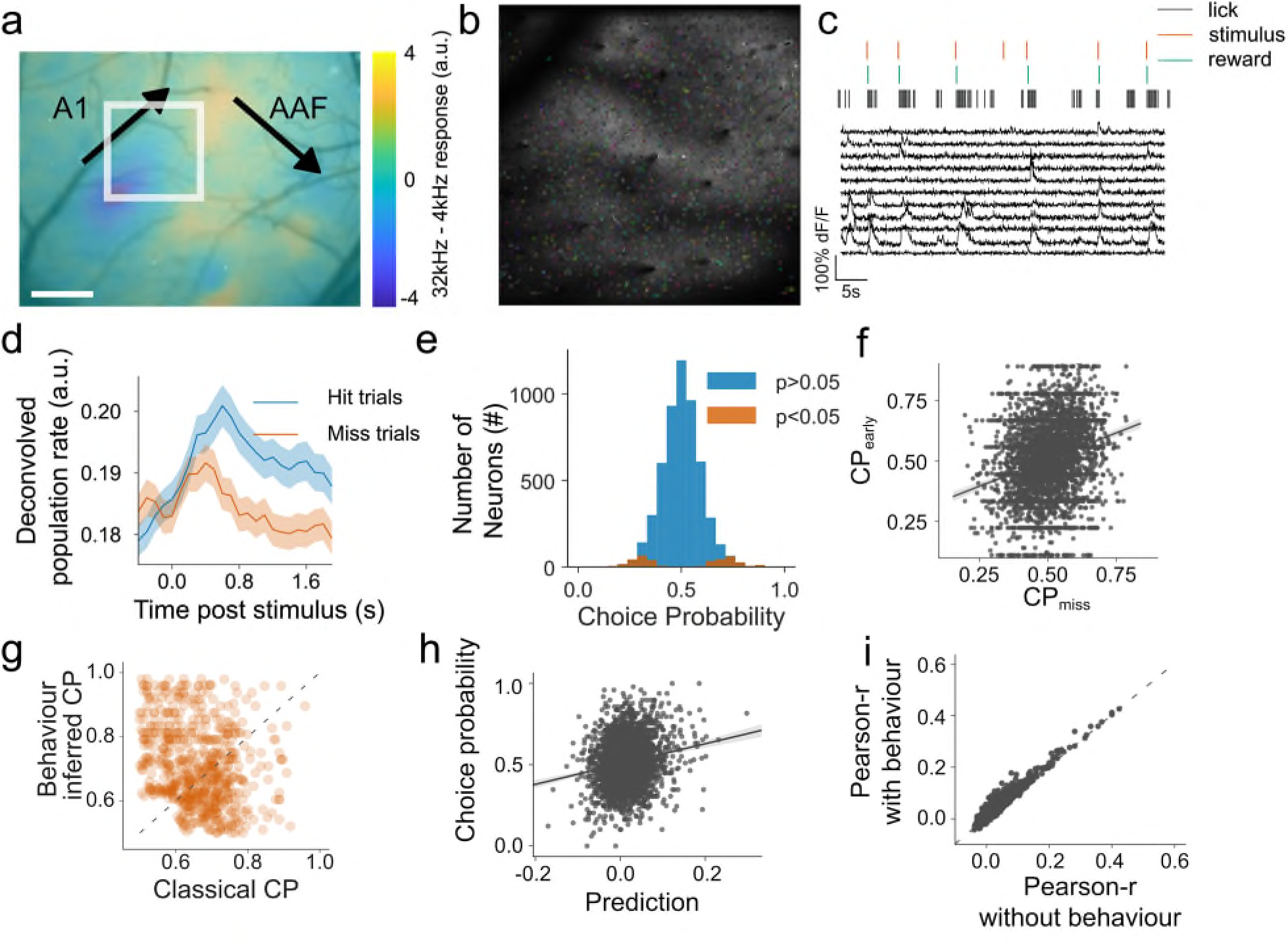
Behaviorally-decoded choices reflect neural activity. **a)** Functional localization of auditory cortical fields using wide-field single photon imaging. Scale bar shows *500μm.* **b)** Example imaging field (~900μm^2^; region in white square in a with regions of interest (n = 976) randomly colored. **c)** Activity of ten neurons from b. **d)** Across the entire population of recorded neurons, we observed significant choice-related activity that emerged shortly after stimulus onset. Shaded regions are ±2 s.e.m. **e)** Distribution of choice probabilities (CPs). Significant CPs (*p* < 0. 05, permutation-test 500 shuffles) were measured in 378 of 5339 neurons (7.1 %). This is a larger subpopulation than would be expected by chance (binomial-test *p* = 2.1 · 10^−119^). **f)** CPs calculated by comparing hit and miss trials and CPs calculated from hit and ‘early hit’ trials are correlated (r = 0.26; *p* = 1.3 · 10^−69^) across neurons. **g)** CPs, plotted here as distance from 0.5, are greater when trial classification is based on model-based decoding rather than *a priori* criteria (paired sample t-test; *p* = 3.6 · 10^−44^). See **supplementary Fig. 6b** for raw CPs. (h) CPs are poorly predicted (meanr^2^ = 1%), on a neuron-by-neuron basis, from neural tuning to behavioral latent states as assessed by fitting a multi-linear regression model. (i) Including behavioral latent states into a linear regression model to predict neural activity significantly improves fit quality (paired sample t-test; p < 1 · 10^−80^).

Given the non-zero false-alarm rates observed in our data, a subset of hit-trials likely occurred as a result of spontaneous behavior, rather than the learned stimulus-response association. In light of the robust choice encoding in A1, we reasoned that, neurally, these trials should more closely resemble miss-trials than hit-trials. If our decoder is able to correctly reclassify those hit-trials on which licking was spontaneous, we should observe larger CPs. Consistent with this expectation, we found that CPs were indeed larger when calculated based on decoded causes of behavior (mean = 0.71; s.e.m=0.005), than on *a priori* criteria (mean = 0.67; s.e.m = 0.0034), i.e. defining all trials with licking in a window 150-600 ms after the stimulus and no prestimulus licking as hit trials (**Fig 3g., Supplementary Fig. 7b**). This suggests that model-based decoding of video data can provide a more accurate readout of behavior than readouts based on *a priori* definitions imposed by the task structure.

Finally, we sought to use the behavioral models to further clarify the relationship between neural encoding of movement-related and choice-related variables. To relate neural activity to these variables, we fitted a linear model that attempts to explain neurons’ frame-by-frame activity using experimental variables as well as behavioral latent-states. This approach allowed us to dissociate movement- and decision-related influences on neural activity, as during the inter-trial interval movement and decisions are decoupled. Fitting these models to the activity of each neuron thus yielded parameters quantifying how the activity of a given neuron covaries with the animal’s behavior. To further examine whether movement-related influences on neural activity underlie CPs, we attempted to predict neurons’ CPs from these parameters. We found that the relationship between a neuron’s activity and behavioral latent states was poorly predictive of its CP (**Fig. 3h**). Together with the behavioral controls (**Fig. 3f**), these findings strongly suggest that neural tuning to motor variables does not underlie choice-related activity in A1.

Recent work has demonstrated that animals’ movements are predictive of neural activity across cortical regions, including sensory cortex^19^. Consistent with this result, we were better able to predict neural activity using both behavioral latent states and experimental variables as regressors, than experimental variables alone (**Fig. 3i**). However, this could either reflect genuine neural tuning to motor output or be mediated via effects of internal variables on both neural activity and motor output. The comprehensive representations learned by the BAE allowed us to differentiate these two possibilities by quantifying how well A1 population activity predicts animals’ movements. If neurons in A1 are truly tuned to motor output, we should be able to accurately reconstruct behavioral latent states from the measured neural activity. Contrary to this prediction, we were poorly able to predict behavioral latent states from neural activity (mean *r*^2^ = 3%; range 1%-5%). These findings strongly argue that motor output has, at most, a small effect on auditory cortical activity and that correlations between the two are likely mediated by variables such as an animal’s decision that affect both movement and neural activity.

In summary, our novel class of Bayesian model enables comprehensive and interpretable quantification of momentary behavior. Application of this model demonstrated robust encoding of cognitive variables in animals’ behavior and enabled us to disentangle neural encoding of cognitive and motor variables. We constructed model-based decoders whose application provided sub-second accounts of behavior which more accurately reflected neural activity than behavioral readouts imposed by task structure. Combined with recent methods for pose estimation^20^, we envision that our approach will be able to extract simple readouts of complex behavior. Finally, while we have deployed our model in the context of a sensory decision-making task, these methods should be broadly applicable to both basic and clinical research seeking to relate neural activity, computation and behavior.

## Acknowledgements

We would like to thank all members of the Auditory Neuroscience Group for comments and feedback on the work. In particular we would like to thank Samuel Picard and Michael Lohse for valuable comments on the manuscript and critical discussion. We would further like to thank Lydia Oikonomidis, James Phillips and Martin Kahn for comments on the manuscript and critical discussion of the project. This work was supported by a Wellcome Trust PhD studentship (WT102373/Z/13/Z) to YW and a Wellcome Principal Research Fellowship (WT07650AIA, WT108369/Z/2015/Z) to AJK.

## Author Contributions

Y.W. conceived the study and the model. Y.W. and J.C.D. designed the experiments. Y.W. and J.C.D. performed surgeries. Y.W. performed experiments. Y.W. analysed the data. A.J.K. provided infrastructure and resources. A.J.K. and J.C.D. supervised the project. Y.W., A.J.K. and J.C.D. wrote the manuscript.

## Competing Interests

The authors declare no competing interests.

## Supplementary Figures

**Supplementary Figure 1.**
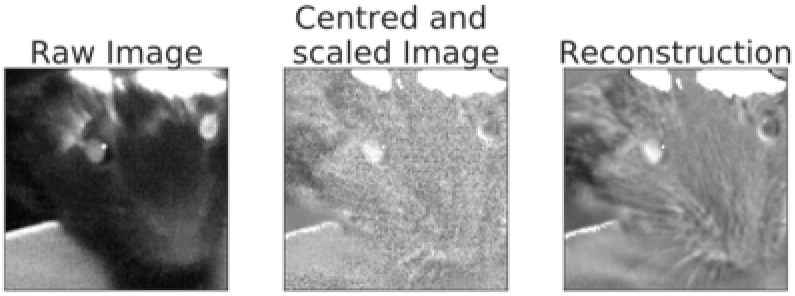
Visualization of reconstructions from the latent space. Example of a video frame in its raw and preprocessed form as well as its reconstruction. In the preprocessing step, each pixel of video data had its mean subtracted and was divided by its variance.

**Supplementary Figure 2.**
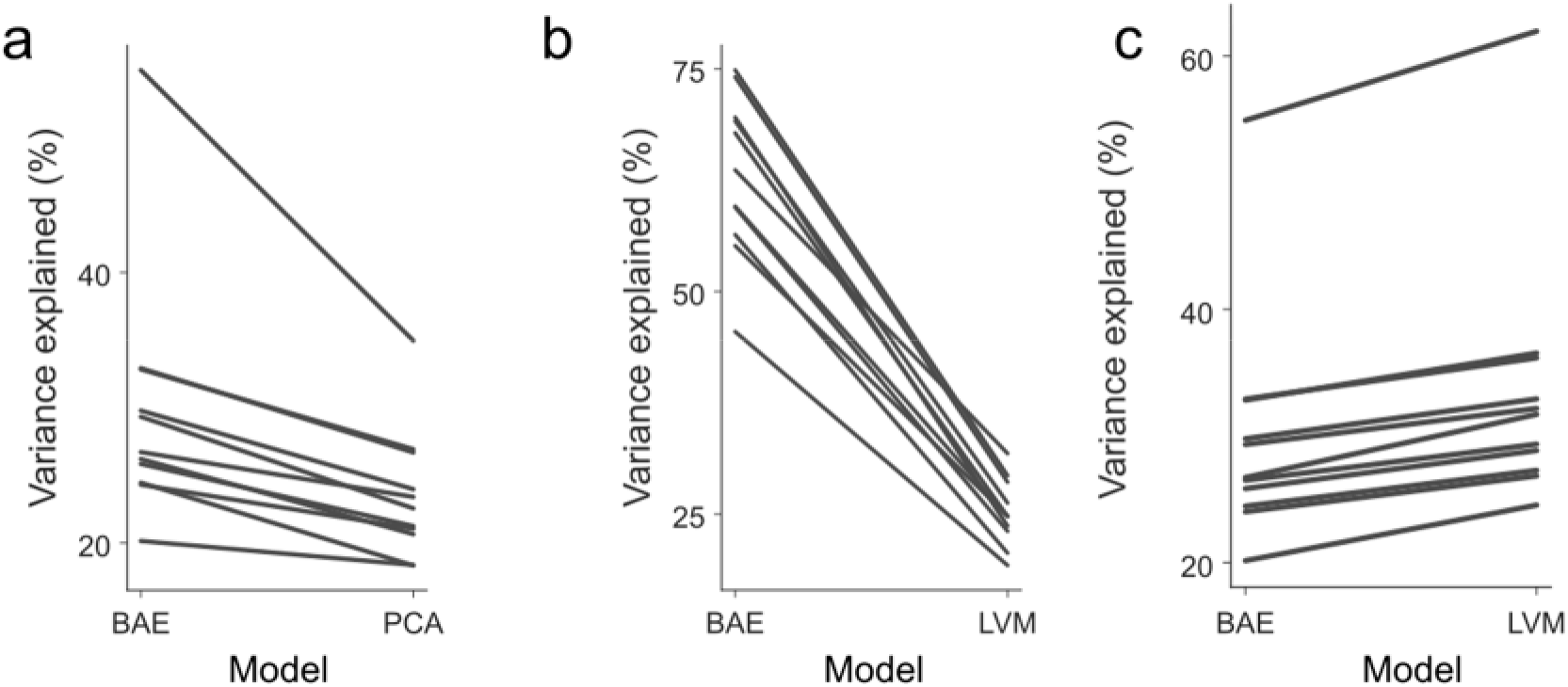
Quantitative analysis of pixel-space reconstructions of video data by various models. **a)** Pairwise comparison of reconstructions of the video data by BAE and PCA. For BAE reconstructions shown here, we performed one full pass through the model, using the recognition model to obtain latent-states and the generative model to obtain pixel-space reconstructions. Each line represents a single session. In all cases, BAE outperforms PCA (paired t-test; p=0.0002). **b)** To assess how well latent states can be predicted from experimental variables we compared the ability of the BAE and LVM (**Fig 1b**) to predict behavioral latent states. The BAE out performed the LVM in all sessions (paired t-test; p=3.5 · 10^−10^), demonstrating enhanced, linear predictability of latent-states as a result of the augmentation of the model’s cost function. **c)** Pixel-space reconstructions, created by a full pass of the video data through the LVM (i.e. video data are passed through the encoder network to generate latent variables, which are then passed to the decoder network, reconstructing the full images) are better than BAE (Paired t-test; p=1.7 · 10^−7^).

**Supplementary Figure 3.**
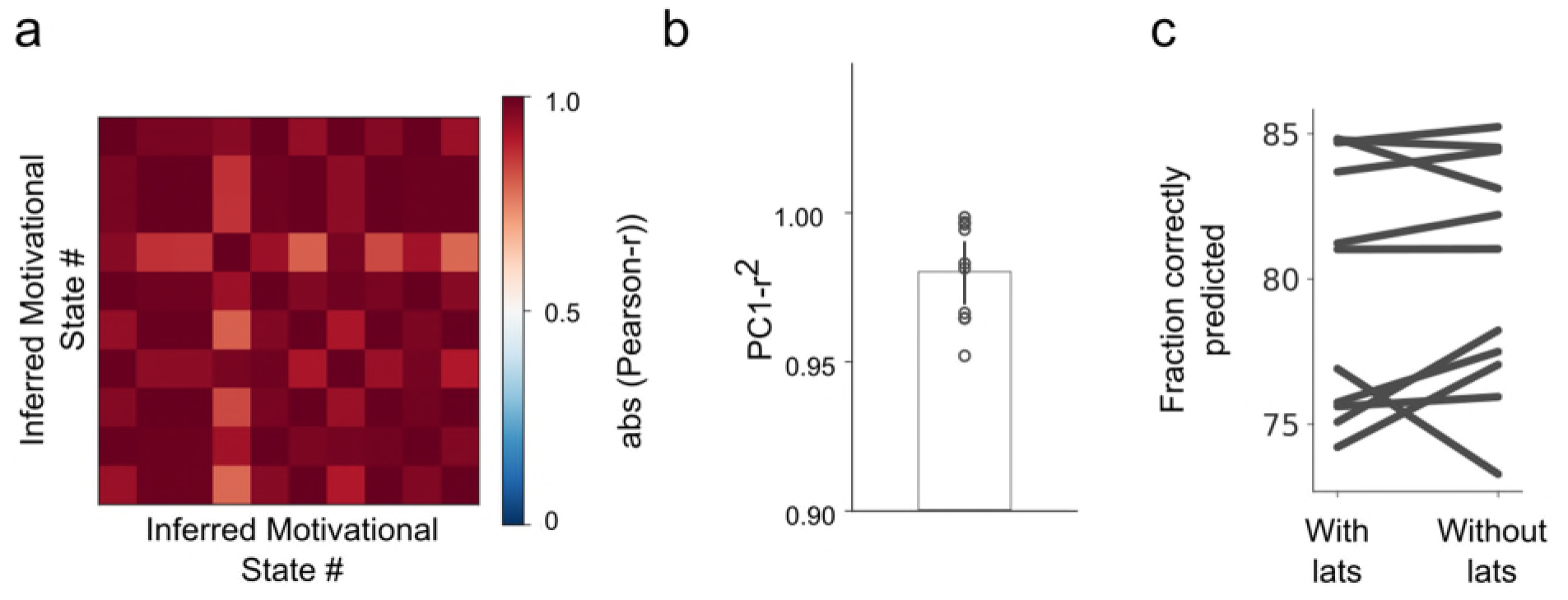
Further analysis of behavioral correlates of cognitive variables. **a)** Analysis of the encoding model from an example session, which shows that motivational state explains variance not accounted for by licking, suggested that an animal’s motivational state is externalized in behavior (**Fig 2a,b**). However, there is a chance that the encoded quantity may not actually reflect motivation, but changes in posture that are unrelated to the animal’s motivational state. Motivation, in the context of our behavioral task, may be measured along a one-dimensional continuum, that is to say that at each point in time animals have a certain level of motivation. Therefore, if the measured quantity truly reflects motivation, we reasoned that different parts of the animal’s posture, reflected in the ten behavioral latent-states, should change in a coordinated fashion. In contrast to this, if the measured quantity is just related to slow changes in posture, there is no *a priori* reason that the different behavioral latent states should change in a correlated fashion. To distinguish these possibilities we calculated the weighted sum of motivation regressors for each latent variable. Regressors were weighted by the values of fitted regression parameters for each latent variable. We refer to this sum as the inferred motivational state. We then measured the correlation between the inferred motivational states fitted to each latent state. Shown is an example correlation matrix, constructed by cross-correlating the inferred motivational states for each latent variable.This example illustrates that inferred motivational states, fitted to each behavioral latent-state independently, are highly correlated, consistent with the hypothesis that the extracted variable is related to the animals’ motivational state rather than arising from spurious changes in posture. **b)** To quantify the extent to which the motivational state variables may be described by a one-dimensional quantity, we performed principal component analysis and quantified the variance explained by the first principal component. We found that in all sessions a single principal component captured more than 95% of the variance across motivational variables. **c)** Analysis of encoding model parameters suggested that attention was not expressed in animal’s behavior. To further test this, we performed a logistic regression analysis and tried to predict trial-by-trial decisions, asking whether knowledge of latent-states preceding stimulus onset helped us in doing so. We compared performance of a baseline model to performance of an extended model that included the latent-states preceding stimulus onset. The baseline model included the intensity of the presented stimulus and whether the previous trial was a hit-or miss-trial. Expanding this model by including behavioral latent states preceding stimulus presentation did not improve the model’s ability to predict whether a given trial is a hit-or miss-trial (paired sample t-test; *p* = 0.32). These results bolster the conclusion that attention is not encoded in the animals’ behavior preceding stimulus onset.

**Supplementary Figure 4.**
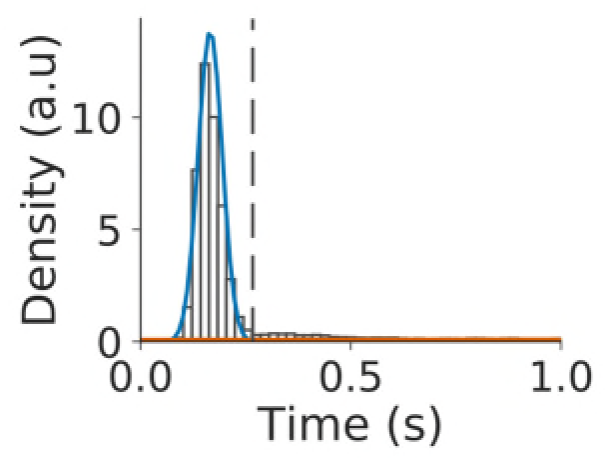
Mouse licking behavior is organized into bouts. Distribution of inter-lick intervals across all sessions and animals (white histogram bars). Gaussians fitted to intra-bout inter-lick intervals (blue curve) and between-bout inter-lick intervals (orange curve) overlaid, together with the optimal separation boundary (dashed vertical gray line).

**Supplementary Figure 5.**
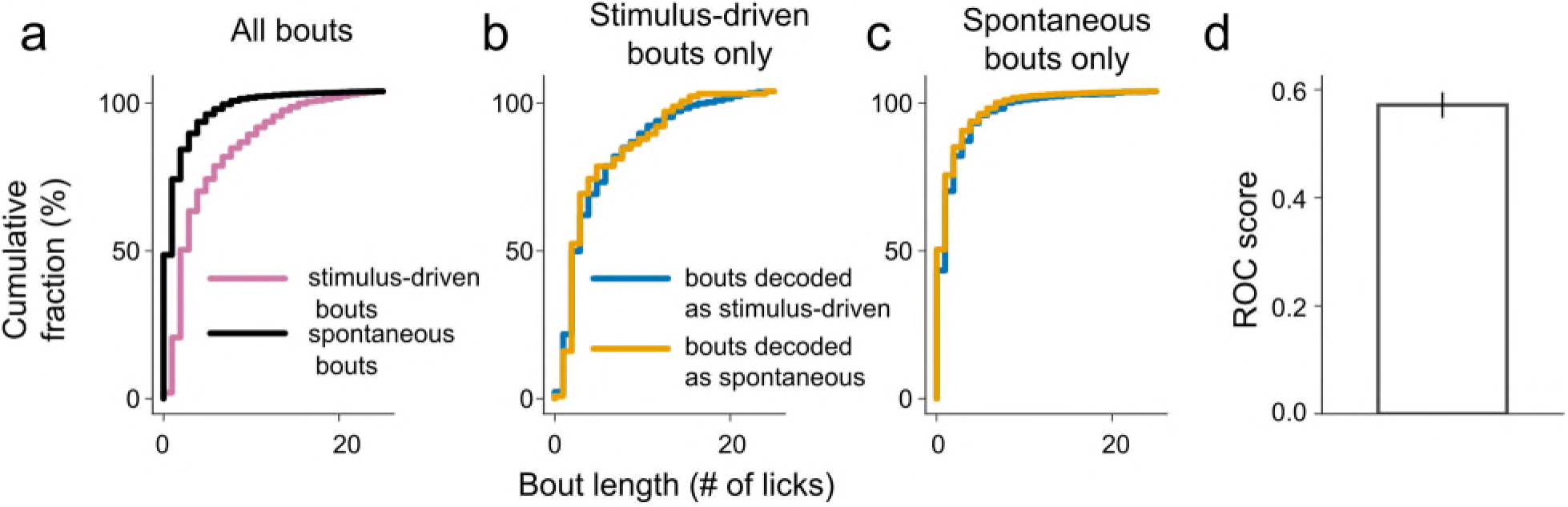
Excluding motor preparation and time as bases for classifying behavior. **a)** Significant differences in bout lengths (quantified in terms of number of licks in a bout) exist between stimulus-driven and spontaneous bouts. Therefore, stimulus-driven and spontaneous bouts could be associated with differences in motor preparation that the decoder might be able to exploit for its classification. **b)** Partitioning of only stimulus-evoked bouts according to decoder classification reveals no differences in bout length as a function of the decoder’s classification. **c)** Partitioning of only spontaneous bouts according to decoder classification also revealed no difference in bout length as a function of the decoder’s classification. This suggests that decoder performance is not driven by potential differences in motor preparation between short and long lick bouts. **d)** To estimate the extent to which the decoder relies on differences in bout length to perform classification, we measured how well bout length could predict decoding performance. To do so, we computed the area under the receiver operating characteristic curve (mean=0.56; s.e.m=0.01) and found that bout length was a poor predictor of the decoder’s decision.

**Supplementary Figure 6.**
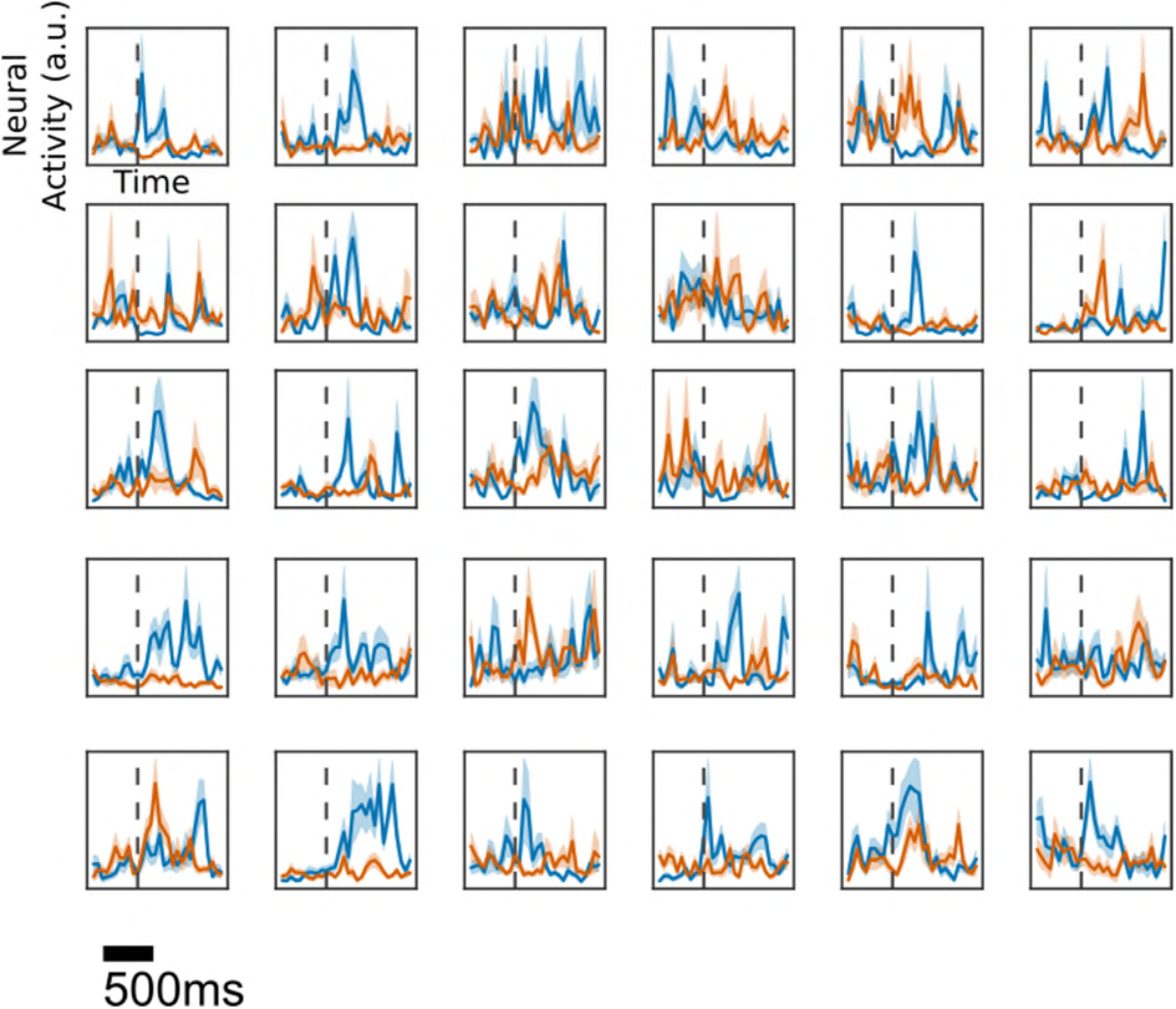
Representative examples of neurons with significant choice probabilities. Each panel shows the average activity (mean±s.e.m) of a single neuron in a window surrounding stimulus onset (dashed vertical line). The y-axis of each panel is normalized to show the full dynamic range of each neuron. Blue curves show mean activity during hit-trials; orange curves show mean activity during miss-trials. Examples shown are taken from all animals.

**Supplementary Figure 7.**
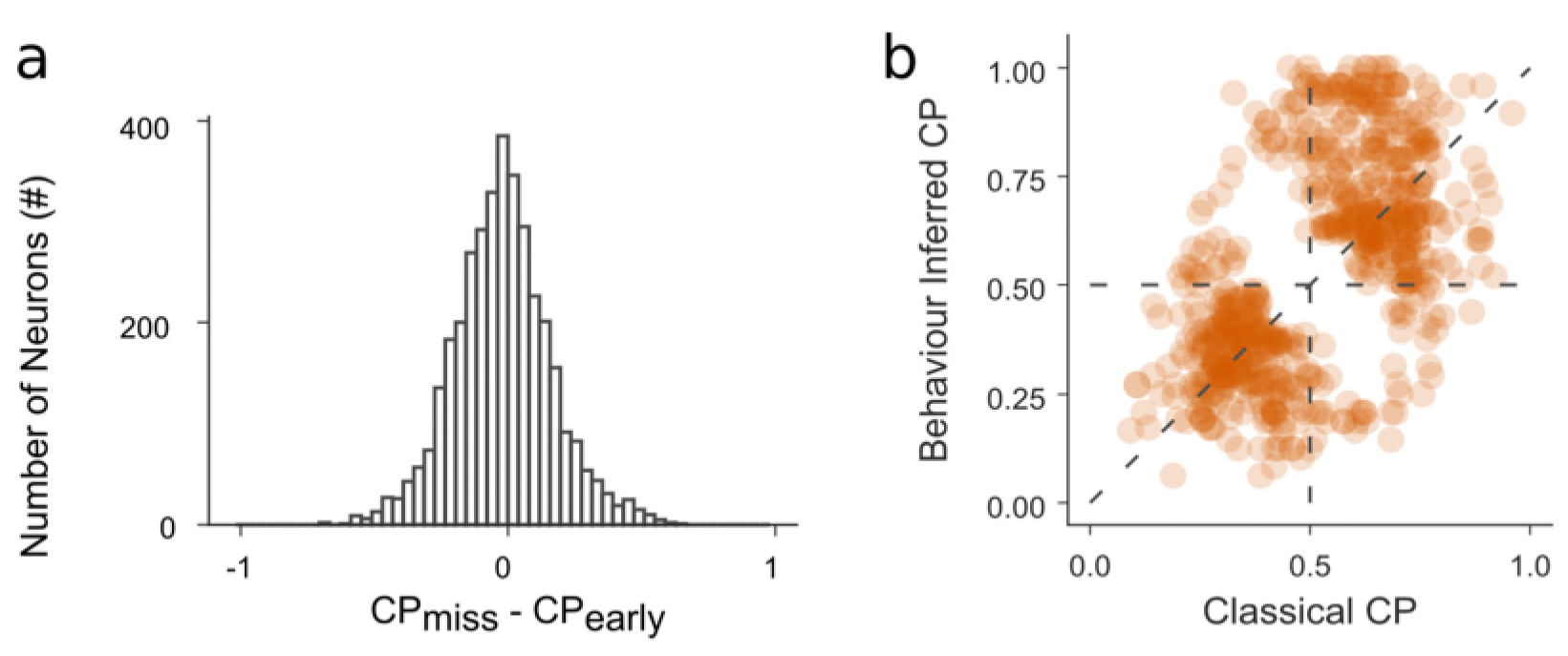
Further analysis of choice-related activity. **a)** Choice probabilities calculated by comparing hit and miss trials (CP_miss_) and choice probabilities computed by comparing hit vs early hit trials (CP_early_) are not significantly different in magnitude (paired-sample t-test *p* = 0.68). **b)** Full distribution of classical versus behavior inferred choice probabilities.

**Supplementary Video 1**. Example pre-processed video and associated reconstructions using the BAE. Latent states were estimated using the recognition model.

**Supplementary Video 2**. Estimation, via the BAE, of the mean video sequence preceding stimulus-driven and spontaneous lick bouts, respectively. Estimation is based on data from one example session. These pre-lick bout sequences were estimated by reconstructing latent states using the behavioral encoding model and projecting these latent states into pixel space using the generative model.

**Supplementary Video 3**. Example sets of video sequences preceding stimulus-driven and spontaneous lick bouts from a single session. Data shown in video are temporally counterbalanced such that simultaneously shown clips are close in time. Data are from the same session as Supplementary Video 2.

## Methods

### Animals

All experiments were approved by the local ethical review committee at the University of Oxford and licensed by the UK Home Office. One female C57BL/6NTac.*Cdh23753A>G* (Harlan Laboratories, UK) mice^23^, 3 female (C57B6.129S-Gt(ROSA)26Sortm95.1(CAG-GCaMP6f)Hze [Jax: 024105] x C57B6.Cg-Tg(Camk2a-cre)T29-1Stl/J [Jax:005359]), one male *(Igs7tm93.1(tetO-GCaMP6f)Hze* Tg(Camk2a-tTA)1Mmay/J [Jax: 024108] x Rbp4_KL100-Cre, MMRRC: 037128; Gerfen et al., 2013) and one male Rbp4-cre mouse were used for behavioral experiments. Neural data were obtained from the three (C57B6.129S-Gt(ROSA)26Sortm95.1(CAG-GCaMP6f)Hze x C57B6.Cg-Tg(Camk2a-cre)T29-1Stl/J) mice. All experiments were performed before mice reached 12 weeks of age, preceding the onset of age-related sensorineural hearing loss in C57BL/6J strains^21,22^.

### Click detection task

Three days before mice commenced behavioral training, we started restricting their access to water and acclimatising them to handling and head-fixation. Throughout the training and testing period the mice’ body weight remained above 80% of their pre-restriction body weight. Mice were trained daily to lick in response to a 0.05-ms biphasic click stimulus presented at 80 dB SPL. There were two types of trials: stimulus trials (80 dB SPL click; water reward for licking) and catch trials (no stimulus; no reward for licking). These were randomly interleaved at an inter-trial interval drawn from a uniform distribution between 6s and 12s. If mice licked during a 1.5 s window following onset of the stimulus, a water drop (2 μl) was delivered immediately. Once mice reached high performance levels (> 80 % correct on stimulus trials), which took 2-5 sessions, they were moved to the testing phase in which stimuli were presented at different intensities. Stimuli were randomly interleaved and presented over a maximum range of 38 dB SPL to 80 dB SPL (3-dB steps). The range of stimulus levels presented in a given session was, in some cases, adjusted according to the animals’ sensitivity. Behavioral data were acquired in blocks lasting between 7 and 30 minutes. Typical sessions lasted approximately forty minutes during which mice performed approximately 250 trials.

Data were excluded, in a block-wise manner according to several criteria. Firstly, mice needed to have undergone at least two testing sessions prior to the sessions considered for inclusion. Secondly, to be able to reliably identify stimulus-driven bouts, we required hit-rates for the loudest stimuli to exceed 95%. Finally, to be able to reliably identify hit-trials as being stimulus driven, we required false-alarm rates to be below 45%. Of the 12 sessions (two per mouse) passing these criteria, one had to be excluded because of video frames missing as a result of camera failure.

### Apparatus

The behavioral apparatus was controlled from a computer running Windows 7 using MATLAB (Mathworks) interfaced with a National Instruments board (NI-DAQ USB-6008) for data acquisition. Stimuli were presented using MATLAB 2016a (Mathworks) running psychtoolbox. Stimuli were digital-to analog converted using a commercial soundcard (ASUS Xonar-U7), amplified (Portable Ultrasonic Power Amplified; Avisoft Bioacoustics) and played through a free-field electrostatic speaker (Vifa; Avisoft Bioacoustics), positioned approximately 15 cm in front of the mouse’s snout.

Stimuli were calibrated using an M500 microphone (Pettersson), which was itself referenced to a sound-level calibrator (Iso-Tech SLC-1356). Click volumes were calibrated by integrating the recorded RMS of clicks over the mouse hearing range (1-100kHz) and comparing it to the RMS of stimuli from the reference sound-level calibrator.

Video frame acquisition was triggered by the frame clock of the two-photon microscope, such that one video frame was acquired for every two microscope frames, resulting in an acquisition rate of ~13 Hz at a resolution of 640 x 480 pixels. The camera, a DMK23UV024 (The Imaging Source) mounted with a M5018-MP2 (Computar) lens, was positioned approximately 30 cm in front of and 30 cm above the behavior apparatus, aligned to have the mouse’s face and most of its body in the field of view. Regions of interest showing the mouse’s face (Supplementary Fig. 1) were drawn manually (approximately 150 x 150 pixels in size) on each dataset. These regions of interest were used for further analysis.

### Widefield calcium imaging

The widefield imaging system consisted of a 470nm LED (M470L3, Thorlabs), a digital camera (340M-GE, Thorlabs) and a 2X objective (TL2X-SAP, Thorlabs) mounted on a Thorlabs Bergamo II microscope body. Images were acquired at a rate of 10 Hz and a resolution of 96 by 128 pixels using ThorCam (Thorlabs) software. Sound waveforms were generated in LabView (National Instruments) and presented on the same hardware as described above. For the frequency mapping of auditory cortical fields we presented 500 ms long sinusoidally amplitude modulated (SAM) tones with a modulation frequency and depth of 10 Hz and 100%, respectively. Each map was based on the responses to 15 repeats of one low carrier frequency (4 kHz or 5.04 kHz) and 15 repeats of one high carrier frequency (25.4 kHz or 32 kHz) SAM tone, presented at either 55 dB SPL or 65 dB SPL and at a rate of 0.33Hz. Frequency maps (Fig. 3a) were generated by calculating the average response (mean signal intensity in a 1-s window following sound onset minus mean signal in a 1-s window preceding sound onset) to the low-frequency and high-frequency stimulus, subtracting one from the other, color-coding the resulting image and superimposing it on a grayscale image of the bloodvessel pattern.

### Two-photon data acquisition

Two photon imaging was performed as described previously^24^. Briefly, image acquisition was carried out using a commercially available two-photon laser-scanning system (B-Scope; Thorlabs). A SpectraPhysics Mai-Tai eHP laser fitted with a DeepSee prechirp unit (70fs pulse width, 80MHz repetition rate) provided the laser beam for two photon excitation. The beam was directed into a Conoptics modulator and then through the objective (16x 0.8NA water immersion objective; Nikon). The beam was scanned across the brain using an 8-kHz resonance scanner (X) and a galvanometric mirror (Y). The resonance scanner was used in bidirectional mode, enabling acquisition of 512 x 512 pixels at a frame-rate of approximately 26 Hz. Emitted photons were filtered (525/50) and collected and amplified by GaAsP photomultiplier tubes (Hamamatsu). ScanImage was used to acquire data and control the microscope. All imaging was done between 150 and *250μm* below the cortical surface.

### Latent variable model

The mathematics underlying variational autoencoders^10,11^, on which our models are based, has been covered in great detail elsewhere (see e.g. Doersch, 2016^25^ for a tutorial) so we will give only a brief summary here. Given some observed high-dimensional series of pixel intensities (i.e. video data) *X,* we seek to explain variation in *X* by assuming that some low-dimensional underlying latent variables, *z*, give rise to the data. Ideally, the quantity we would seek to maximize when fitting the model is thus*P(X)*, the probability of the data. We can relate zto *P(X)* mathematically by conditioning:

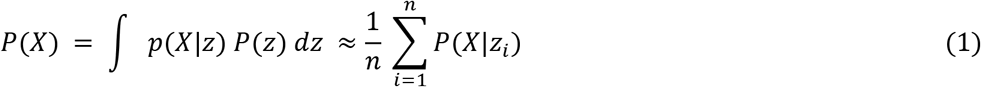

where we note that any integral can be approximated by a finite sum over samples of *z_i_*. This formulation has the important property that by specifying the functional form of *p(X|z)* and a method of sampling *z_i_* we can evaluate *P(X)* and hence quantify the performance of the model. For analytical tractability and ease of sampling, we assert that *P(z)* is a Gaussian distribution with 0 mean and diagonal, unit covariance.

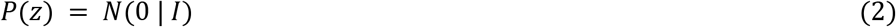

Based on the continuous values of pixel intensities, we further specify *P(X_i_|z_i_*) to be a normal distribution:

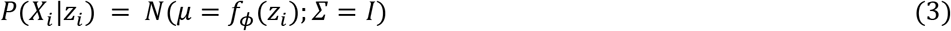

where *f_φ_*(*z*) is a deterministic function, with parameters *φ*, that map latent variables, *z*, into pixel space. In practice, we implement *f_φ_*(*z*) as a multi-layer neural network.

However, with high-dimensional data, naive sampling approaches are inefficient to the point of intractability because for most values of *z_i_*, *p*(*X_i_* |*z_i_*) ≈ 0. To enable efficient sampling, allowing us to tractably approximate the above integral, we construct an auxiliary distribution *Q*(*z_i_*|*X_i_*) which enables us to draw samples from *P*(*z_i_*) such that the sampled *z_i_* are likely to give rise to *X_i_*. In practice, we assume that

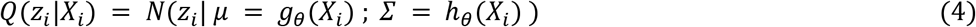

where *g* and *h* are deterministic functions of *X*, parameterised by *θ,* which are implemented by a deep neural network. However, naively sampling *Q*(*z|X*),rather than *P*(*z*), to evaluate *P(X)* will result in biased estimates. To circumvent these issues we apply standard identities from the Variational Bayesian literature^7^ to derive:

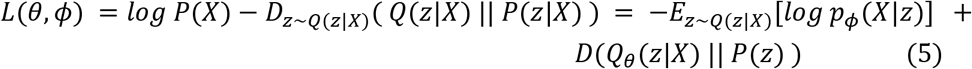

where *D*(*p||q*) denotes the KL-Divergence (a measure of difference between probability distributions) between *p* and *q.* The left hand side of this equation is the quantity we seek to maximize. Doing so maximizes the likelihood of the data *P(X)* while minimizing the difference between our approximation of *Q(z|X)* and the true *P(z*|X). Since both *Q_θ_*(*z|X*) and *P(z)* are Gaussian, this divergence has a closed form solution. Similarly, we can arrive at a computationally tractable form of the expectation *E_z~Q(z|X)_*[·] by using a single sample from *Q*(*z|X)* to make the approximation. Furthermore, tractable derivatives of this cost function are available^10,11^.

We extend this model to encourage learning of interpretable latent representations. We achieved this by adding an additional term to the cost function. Specifically, we fitted a behavioral encoding model (see *Behavioral encoding model* for details), mapping from task variables to the latent variables *z* using a linear regression model with parameters *β.* We augment the cost function with the error term of this regression model to obtain a more interpretable model in which the values of latent variables *z* are linearly predictable from variables of interest.

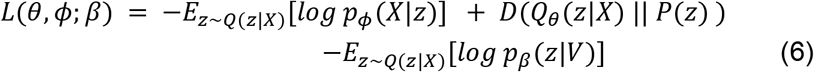

Importantly, the prior on the latent space acts to regularize the latent parameters preventing overfitting. Additionally, our behavioral encoding model only biases the learning of weights, it does not bias the inferred latent representation.

### Data analysis

Data were analysed in Matlab and Python 3.6.2 augmented with standard libraries for scientific computing^26–31^. Unless stated otherwise, standard algorithms (e.g. principal component analysis) are implemented using reference implementations from these libraries. A reference implementation of the behavioral autoencoder, together with an example video dataset is available for use and alteration at www.github.com/yves-weissenberger/bae.

All statistical tests were, unless otherwise stated, implemented using reference implementations in standard libraries for scientific computing in Python. All statistical tests were two-tailed.

### Model implementation

The hierarchical Bayesian model is implemented using the Python library *Tensorflow*^32^. The model is comprised of two sequential networks termed recognition model and generative model, respectively. All neural activation functions were rectified-linear unless otherwise stated.

The recognition model is a four-layer network. The first two layers are comprised of convolutional units (256 and 128 units), and kernel sizes three and five pixels, respectively. In both cases, the stride of kernels was set to two pixels. These layers were followed by a fully connected layer with 100 units and a final bipartite layer comprised of 10 linear and 10 softplus units, mapping to the mean and covariance of the latent space, respectively.

The decoder network consisted of two fully connected layers with 100 and 500 units, respectively, followed by a final fully connected linear layer mapping the previous layers’ output into pixel space. Our network was trained using a 60/20/20 train/validation/test split. To optimize the cost function we used AdamOptimizer^33^ with the learning rate set to 0.005. Hyperparameters were, once heuristically optimized using a separate dataset not included in this report, held fixed for all analyses reported here.

### Lick bout analysis

To separate licks into bouts, we fitted a two-component Gaussian Mixture Model (implemented by the *GaussianMixture* class of the scikit-learn library) to the inter-lick interval (ILI) distribution of all mice. We thereby separated the ILI distribution into two components which we interpreted as corresponding to within bout ILIs and across bout ILIs. In doing so, we determined the optimal separation window for dividing licks into bouts as the point at which the probability of the fitted Gaussian with the larger mean exceeded that of the smaller one. Doing so, we found that a window of ~266ms provided the optimal separation window for differentiating within-bout licks from across-bout licks.

### Behavioral encoding model

Our behavioral encoding model was a linear-regression model mapping from the set of observed and hidden variables*V*to inferred latent-states *z_i_* using parameters *β*. The set of observed variables we used comprised licks, rewards, lick-bout initiations (defined as the first lick in a bout of licks) and sound stimuli. The timestamps of each of these observed event types were discretized to construct a set of *T* × 1 vectors (where *T* is the length of the session), either set to 1 on the camera frame at which the event occurred (click, reward) or two frames preceding an event (lick-bout initiation, lick), as these movements will be initiated before a lick is completed, and 0 everywhere else. In the case of the clicks, we also analyzed the data after scaling entries in the vector according to sound level, but this made no qualitative or quantitative difference (data not shown).

The set of hidden variables was comprised of decision basis, attention and motivational state. Decision basis was a *T* × 2 binary vector whose first and second columns signified whether a stimulus-driven or spontaneous lick-bout occurred, respectively. An entry in the first column was set to a value of 1 at five frames (~380 ms) preceding the onset of a lick-bout if a stimulus preceded the lick-bout within a ~600 ms window (this window represents the 70th percentile of the across-animal reaction time distribution). Analogously, an element was set to 1 in the second column if no stimulus preceded the bout and the bout was initiated outside the peri-stimulus period. This period was defined as the period from ~150 ms prior to onset of the stimulus to ~1.5 s following the onset of the stimulus.

Attention was a *T* × 2 binary vector whose first column signified that the animal was attentive. We reasoned that detection of particularly loud stimuli was not affected by attention and therefore did not include these in this analysis. An element in the first column was set to 1 at five frames preceding the onset of a stimulus if that stimulus was presented at a low intensity (average hit-rate at that intensity <75%) and the trial was a hit trial. Analogously, an element in the second column was set to 1 on miss trials.

Motivational state was a *T* × 5 continuously valued vector approximating the extent of reward seeking. We constructed each row of this matrix by convolving the vector of licks with a Gaussian distribution. We derived this definition of motivational state based on recent work demonstrating that in head-fixed mice, increased motivation is associated with increased baseline lick rates^34^. The Gaussian for each row had a different standard deviation reflecting our *a priori* uncertainty about the timescales of motivational fluctuations. The standard deviations ranged from ~2.5 s to ~40 s multiplied in powers of two.

We additionally included a set of time regressors, a *T* × 10 vector, where each row is a continuous low frequency oscillation, to account for slow drifts in posture over time. The period of these oscillations ranged from ~1450 s to ~2150 s. To enable events to affect latent-states at future time points, all the above vectors (with the exception of motivational-state and time) were multiplied with a Toeplitz matrix giving rise to a series of lagging regressors extending 5 frames into the future.

The Design Matrix 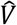 was then constructed by concatenating these vectors together with an offset term yielding the following regression model

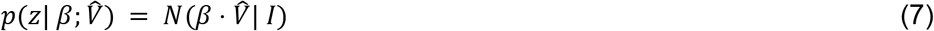

where

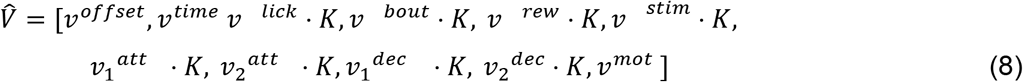

Linear models were regularized using an L2-penalty term. Fitting, as well as regularization parameter selection was implemented using the scikit-learn function *RidgeCV.* Fit quality estimation was performed using repeated, nested K-fold cross validation (five folds; four repeats). In the inner K-fold loop (five folds), the training data were used for fitting and hyperparameter selection, while in an outer loop fit quality was assessed using the held-out data.

### Analysis of behavioral-encoding model parameters

To determine the importance of each regressor in the behavioral encoding model, we performed two complementary analyses to bound the extent of their encoding. This was required because of the collinearity of regressors. To obtain a lower bound on strength of encoding, we quantified the effect of excluding subsets of regression parameters, relating to a single experimental variable (e.g. *v*^bout^), on cross-validated fit quality. Secondly, to obtain an upper bound, we included only parameters relating to a single experimental variable in the regression model. Each of these models was fitted to latent-states extracted after the initial, global fitting process. Model performance was estimated, as during initial fitting, using repeated, nested K-fold cross validation (six folds; four repeats). In the inner K-fold loop (five folds), we determined the optimal regularization parameter. In the outer loop, we attempted to assign hit or miss labels to a held-out test set of trials based on fit parameters.

### Logistic-regression analysis of attentional state

To determine whether trial-by-trial attentional states were externalized in behavior, we attempted to use behavioral latent-states preceding stimulus onset to predict whether a given trial was a hit or miss trial. To do so, following fitting of our latent variable model and the determination of behavioral latent-states, we fitted a logistic regression model to subjects’ trial-by-trial choices. Logistic regression was implemented using the *sklearn* function *LogisticRegression* using the Newton Conjugate Gradient solver and an L2 penalty. A reference model included as regressors the level of the presented stimulus and a variable indicating whether the previous trial was a hit- or miss-trial. To determine whether some correlate of attention was externalized in behavior, we compared performance of the reference model to a model which additionally included the behavioral latent states on the ten video frames preceding each stimulus onset as regressors. Model performance was estimated using a repeated, nested K-fold cross validation (six folds; four repeats). Regularization parameters were optimized in an inner K-fold loop (five folds).

### Behavioral decoding dataset

The window for decoding extended 5 video frames backwards from the onset of the lick-bouts. To ensure that lick history did not form the basis of our behavioral decoding, we only selected lick-bouts in which no licks occurred in a ~610 ms window preceding bout-onset. Additionally, to ensure that long-timescale covariation in posture and spontaneous bout-rates do not drive decoder performance (spontaneous bout-rates are typically higher at the beginning of behavioral sessions), spontaneous and stimulus-driven lick-bouts were selected in a temporally counterbalanced fashion. Specifically, for each session, we counted the number of stimulus-driven and spontaneous bouts. We denote the smaller of these two sets the reference set *R*_1_. For each bout in the reference set, we selected the bout in the larger set that was its nearest neighbour, yielding a second set of bouts *R*_2_. The union of these sets (*R*_1_ ∪ *R*_2_) then comprised the decoding dataset. This led to an unbiased selection of spontaneous and stimulus-driven bouts. Decoding performance was similar when the bout distributions were not counterbalanced in this fashion (data not shown). Decoding performance was estimated on a test-set held out during fitting, using repeated, nested K-fold cross validation (five folds; four repeats).

### Model free decoding

Model free decoding was performed using a linear support vector machine whose regularization parameter *C* was determined in an inner cross validation loop, as described above. In addition to determining the optimal regularization parameter, variable selection was performed in the inner loop, whereby the optimal set of timepoints to use for classification was determined by optimizing prediction accuracy on the training set. Classification was implemented by the *sklearn* function SVC.

### Model-based decoding

Decoding was performed using log-likelihood ratios *(LLR)* similarly to Pillow et al^14^. Specifically, for each lick-bout we compared the log-likelihood of the behavioral latent-states preceding the onset of a bout under the assumption that this bout was stimulus-driven, with the log likelihood that the bout was spontaneous:

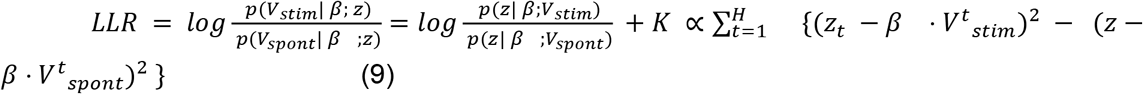

Where *V^t^_stim_* is the design matrix constructed by setting the relevant entry (i.e. five frames preceding bout onset) for stimulus-driven bout to 1 and the entry for spontaneous bout to 0, *V_spon_t* is the reverse, *H* is the analysis horizon and *K* are terms independent of *V.* A log likelihood ratio greater than 0 corresponds to a lick bout that is decoded as being stimulus-driven.

To quantify the accuracy of the decoder we performed a repeated nested, stratified K-fold (six folds; four repeats) cross validation. In an inner K-fold loop (five folds), we determined the optimal regularization parameter for the behavioral encoding model. This means that regularization parameters were only explicitly optimized for encoding, and only implicitly optimized for decoding. Decoding performance was then estimated on the held-out cross validation set comprising equal numbers of stimulus-driven and spontaneous lick-bouts.

Pixel space decoding was performed by projecting latent-space estimates of stimulus-driven (i.e. *β* · *V^t^_stim_*) and spontaneous lick bouts (i.e. *β* · *V^t^_spont_*) back into pixel space using the trained generative model and calculating log likelihood ratios in pixel space.

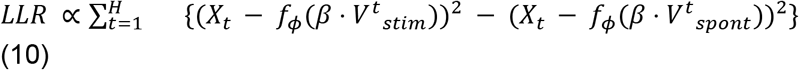

Where *f_φ_*(·) (see equation (3)) is a neural network implementing the generative model, returning the posterior mean in pixel space from some latent value.

### Two-photon data preprocessing

Data preprocessing was performed in Python using the Two-Photon Analysis Toolbox: twoptb (https://yves-weissenberger.github.io/twoptb/). Briefly, data were motion registered using the efficient subpixel registration algorithm. Next, regions of interest (ROIs) were automatically segmented (then manually curated) using a pre-trained supervised algorithm, included in the toolbox, which uses the mean image to identify ROIs. Segmentation was performed in a two-step process where the initial step involved finding seed regions for ROIs using a random-forests classifier. In a second step, a region-growing algorithm was applied to construct ROIs. Traces were extracted as an unweighted average of fluorescence within each region of interest. All traces were neuropil corrected using the fluorescence averaged in a 20 x 20*μm*square surrounding the ROI (empirically determined correction factor: ~0.5). Traces were then baseline corrected using a Kalman-filter based estimate of baseline fluorescence. Finally, spike inference was performed on neuropil corrected traces using the c2s toolbox^35^. To improve temporal resolution, all neural analyses were performed on inferred spike rates.

### Choice probability estimation

For analysis of choice probabilities^12^, we selected equal numbers of hit and miss trials from each stimulus level with hit-rates between 25% and 75%. This was done to maximise data inclusion while preventing variation in sound-evoked activity from dominating the influence of choice. To calculate choice probabilities, we measured the neural response (average neural activity in a 300ms window following stimulus onset) for each trial. We then used the resulting hit and miss trial response distributions to calculate the area under the receiver operating characteristic curve using the *roc_auc_score* function in the *sklearn* package. P-values for choice probabilities were determined by permutation testing using 2000 shuffles.

When calculating choice probability based on behavioral decoding, the subset of hit-trials that were behaviorally decoded as spontaneous were moved from the hit-trial to the miss-trial group. To avoid biased estimates as a result of class imbalances, we calculated choice probability by averaging the mean accuracy for each class (hit and miss). Calculating choice probabilities without such counterbalancing did not qualitatively affect conclusions (data not shown).

### Neural regression model

Regression models fitted to neural activity were identical in implementation to those used in the behavioral encoding model (see above), except for the inclusion of instantaneous (i.e. no time lagged regressors were used) behavioral latent-states as regressors. When neural regression models were fit only to behavioral latent-states and did not include the design matrix used in the behavioral encoding model, results with respect to choice encoding were qualitatively similar (data not shown).

### Choice probability prediction

To assess whether neural choice probabilities (CPs) were related to the covariation of neural activity and movements, we analyzed the parameters of fitted neural regression models. Following the fitting of neural regression models, parameters relating to behavioral latent-states were extracted. We then fitted a multi-linear model, separately to each session, which attempted, on a neuron-by-neuron basis, to predict the neuron’s choice probability from that neuron’s regression model parameters related to behavioral latent-states. We reasoned that if choice probability was explained by neural tuning to motor output, or indeed motion artifacts unaccounted for by image registration, then, across neurons, choice probability should be predictable from neurons’ tuning to behavioral latent states. The multi-linear model was implemented by the *OLS* class from the *statsmodels* library.

